# Thyroid Hormone Dependent Transcriptional Programming by TRβ Requires SWI/SNF Chromatin Remodelers

**DOI:** 10.1101/2021.03.22.436429

**Authors:** Noelle E. Gillis, Joseph R. Boyd, Jennifer A. Tomczak, Seth Frietze, Frances E. Carr

**Author notes:** Co-corresponding Authors Frances E. Carr, PhD, Department of Pharmacology, Larner College of Medicine, University of Vermont, 89 Beaumont Avenue, Burlington VT 05405., Seth Frietze, PhD, Department of Biomedical and Health Sciences, College of Nursing and Health Sciences, University of Vermont, 106 Carrigan Drive, Burlington VT 05405.

## Abstract

Transcriptional regulation in response to thyroid hormone (T_3_) is a dynamic and cell-type specific process that maintains cellular homeostasis and identity in all tissues. However, our understanding of the mechanisms of thyroid hormone receptor (TR) actions at the molecular level are actively being refined. We used an integrated genomics approach to profile and characterize the cistrome of TRβ, map changes in chromatin accessibility, and capture the transcriptomic changes in response to T_3_ in normal thyroid cells. There are significant shifts in TRβ genomic occupancy in response to T_3_, which are associated with differential chromatin accessibility, and differential recruitment of SWI/SNF chromatin remodelers. We further demonstrate selective recruitment of BAF and PBAF SWI/SNF complexes to TRβ binding sites, revealing novel differential functions in regulating chromatin accessibility and gene expression. Our findings highlight three distinct modes of TRβ interaction with chromatin and coordination of coregulator activity.

## INTRODUCTION

Thyroid hormones play key roles in maintaining cell identity, metabolism, and homeostasis in all tissues. Thyroid hormone receptors (TRs) function primarily as transcription factors that regulate broad networks of target genes in response to fluctuations in thyroid hormone (T_3_) levels. Two distinct genes encoding TRα (*THRA*) and TRβ (*THRB*) are expressed in a tissue-specific patterns. TRβ is the predominant TR expressed in the liver, kidney, and thyroid, while TRα is expressed the heart, bone, and brain^1^. The classic bimodal switch model (reviewed in^2,3^) where TR is constitutively bound to chromatin and T_3_ binding promotes dissociation of corepressors and recruitment of coactivators, was the predominant model used to describe transcriptional regulation by TRs for many years. However, a number of studies that point to more nuanced mechanisms have recently been put forth. For example, multiple genome-wide studies of TR binding have shown significant T_3_-dependent recruitment of TRβ to chromatin and *de novo* chromatin remodeling, in opposition to the bimodal switch model^4-6^. Additionally, it has been demonstrated that a large proportion of TRβ binding occurs in distal regulatory elements, and that TRβ coordinates histone acetylation of enhancer regions and higher order chromatin structure^7^. Recruitment of cofactors by TRβ has been recently described as a T_3_-dependent coregulator shift, rather than a complete loss of corepressors and gain of co-activators upon ligand binding^8^. Our data supports a multi-modal regulation model for TRβ interaction with chromatin that integrates each of these concepts.

ATP-dependent chromatin remodeling complexes are known to act as coregulators for nuclear receptor transcriptional regulation and have been implicated in chromatin remodeling specifically by TRβ. SWI/SNF components were found to be associated with NCOR, a well-known co-repressor for TRs and other non-steroidal hormone receptors^9^. This provided indirect evidence that SWI/SNF chromatin remodeling might be required for target gene repression by TRs. We have described an interaction between the SWI/SNF core subunit BRG1 and TRβ for repression the oncogene RUNX2^10^. Recruitment of the SWI/SNF complex to T_3_-activated promoters has been suggested to be dependent upon interactions between SRC and p300, rather than a direct interaction between BRG1 and TR^11^. Direct interaction between BAF57, a key SWI/SNF subunit, and TRβ have been observed at T_3_-responsive promoters^12^. Given the limited studies of SWI/SNF participation in T_3_-regulated gene expression, a specific mechanism for TR interactions with SWI/SNF complexes has yet to be defined.

In this study, we mapped the binding sites of endogenous TRβ in normal thyroid cells and integrated this data with changes in chromatin accessibility to classify three distinct modes of TRβ chromatin interaction and remodeling. However, a majority of TRβ binding sites were found in distal regulatory elements. In order to identify the protein-protein interactions involved in T_3_-dependent transcriptional regulation, we performed a proximity labeling assay followed by mass spectrometry. Some interactions between TRβ and its binding partners are gained or lost with T_3_, but notably the majority do not exhibit a T_3_-dependent switch. Differential enrichment of interactions between TRβ and members of BAF and PBAF SWI/SNF complex subspecies were identified. We further demonstrated that TRβ differentially recruits SWI/SNF complexes to its binding sites. Based on our comprehensive genomic and proteomic analyses, we propose a new model whereby selective recruitment of BAF and PBAF SWI/SNF complexes to TRβ binding sites regulates chromatin accessibility and gene expression.

## RESULTS

### TRβ binds to and remodels chromatin in three distinct modes

We profiled the genomic binding patterns of TRβ in the human normal thyroid epithelial cell line Nthy-ORI using CUT&RUN^13^. This approach allows high-resolution detection of genomic binding sites of low abundance transcription factors compared with ChIP-seq. Replicate CUT&RUN peaks were used to generate high confidence peak sets for TRβ with and without 10nM T_3_ for 6 hours (**Supplemental Figure 1A**). We identified a total of 8,200 TRβ binding sites that we classified into three distinct groups based on response to T_3_ treatment (**Figure 1A**). The largest group is the liganded TRβ binding sites (n = 6,768), which have significantly increased TRβ enrichment when T_3_ is present. Unliganded TRβ binding sites (n = 959) are enriched with TRβ only in the absence of T_3_, and ligand-independent binding sites (n = 473) are detected both in the presence and absence of T_3_. The enrichment profile of TRβ binding at each of the three groups of sites was visualized in a heat map (**Figure 1B**), and in representative genome browser shots (**Figure 1C**). Accordingly, we observed a nearly complete loss of TRβ enrichment at unliganded binding sites with T_3_ treatment, while the enrichment remained stable at ligand-independent binding sites with T_3_ treatment. Liganded binding sites exhibited a significant gain in TRβ enrichment upon T_3_ treatment. The gain in signal at liganded binding sites is consistent with the dynamic assisted loading model^14,15^, the concept whereby nuclear receptors have sparse transient interactions with chromatin and ligand binding increases the residency time to stabilize interactions with response elements.

**Figure 1.**
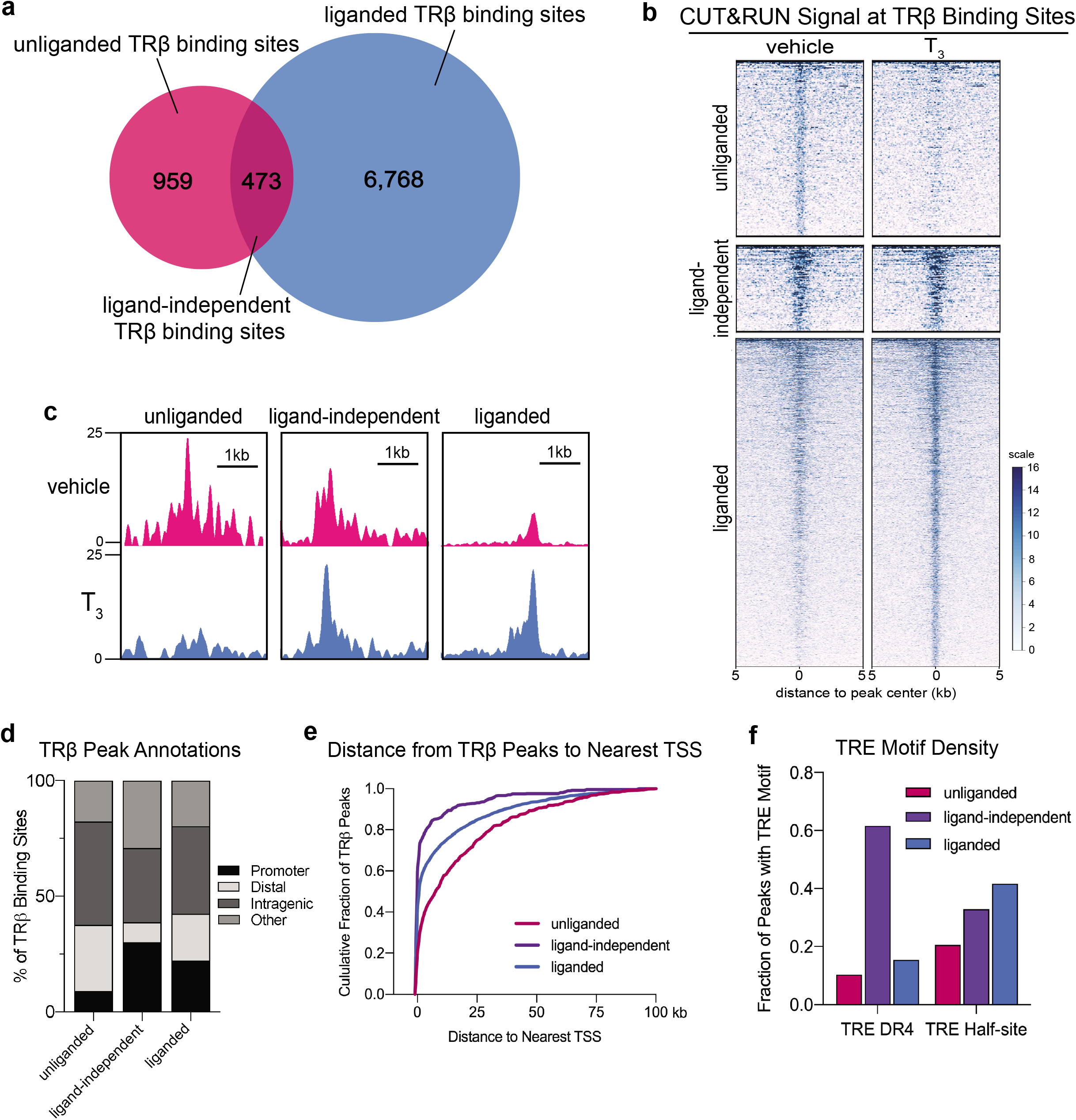
CUT&RUN reveals three distinct modes of TRβ binding in thyroid cells . A. Venn diagram illustrates peak overlap analysis and defines three modes of TRβ occupancy. B. Heatmap demonstrating differentially bound regions classified as unliganded, ligand-independent, or liganded. C. Representative genome browser shots (fold enrichment over lgG) of differentially bound regions. D. Distribution of TRβ binding sites annotated to proximal promoters (< 500bp from TSS), distal regulatory elements (500bp – 10kb from TSS), and intragenic regions. E. Distance from TRβ binding sites to the nearest TSS. F. Fraction of TRβ peaks containing a full-length DR4 thyroid hormone response element (TRE) or a TRE half-site.

Annotation of TRβ binding sites revealed differences in the distribution of TRβ binding relative to transcriptional start sites (TSS) (**Figure 1D**). 30% of ligand-independent sites mapped to proximal promoter regions (within 500bp of TSS), while only 18% of liganded and 9% of unliganded TRβ binding sites mapped to a proximal promoter. Unliganded binding sites were more likely to map to distal regulatory elements (28%), compared to liganded binding sites (19%). Calculating the distance to the nearest TSS of each of these three types of binding sites revealed that ligand-independent binding sites tend to be closer to transcriptional start sites than others (**Figure 1E**). Transcription factor motif analysis revealed that thyroid hormone response elements (TRE) were strongly overrepresented in each of the three groups of TRβ binding sites (**Supplemental Figure 2**). Ligand-independent binding sites have the highest frequency of the full-length direct-repeat palindromic TRE (DR4), while liganded sites have the highest frequency of TRE half-sites.

To map changes in chromatin accessibility in response to T_3_ in Nthy-ORI cells, we performed ATAC-seq after 6 hours (early) and 24 hours (late) hours after treatment with T_3_ (**Supplemental Figure 3**). Differential accessibility analysis revealed a significant increase in the chromatin accessibility with early and late T_3_ treatments (7,754 and 21,678 opened regions, respectively; FDR < 0.05). In contrast, we found that only 107 regions were closed by T_3_ treatment, all of which were closed with early T_3_ treatment (**Figure 2A**). We next examined the chromatin accessibility associated with TRβ binding. Unliganded TRβ binding sites, which are lost upon T_3_ treatment, had relatively little T_3_-induced chromatin accessibility with a modest decrease in accessibility with T_3_ treatment (**Figure 2B**). However, chromatin accessibility increased at liganded binding sites with early T_3_ treatment and further increased with late T_3_ treatment (**Figure 2C**). Ligand-independent binding sites also had significant increases in chromatin accessibility after 6 and 24 hours of T_3_ treatment (**Figure 2D**). To estimate the size of differentially accessible regions of chromatin following T_3_ treatment, we compared the average peak width of differentially accessible ATAC-seq peaks within 5kb of each of the groups of TRβ binding sites (**Figure 2E**). Unliganded TRβ binding sites had the smallest average peak width (190 bp), consistent with the modest changes in accessibility near those sites. Differentially accessible peak width was greatest for ligand-independent TRβ binding sites (355 bp) followed by liganded sites (253 bp). This suggests that remodeling around liganded binding TRβ binding sites is more focused while remodeling around ligand-independent TRβ binding sites stretches across broader regions.

**Figure 2.**
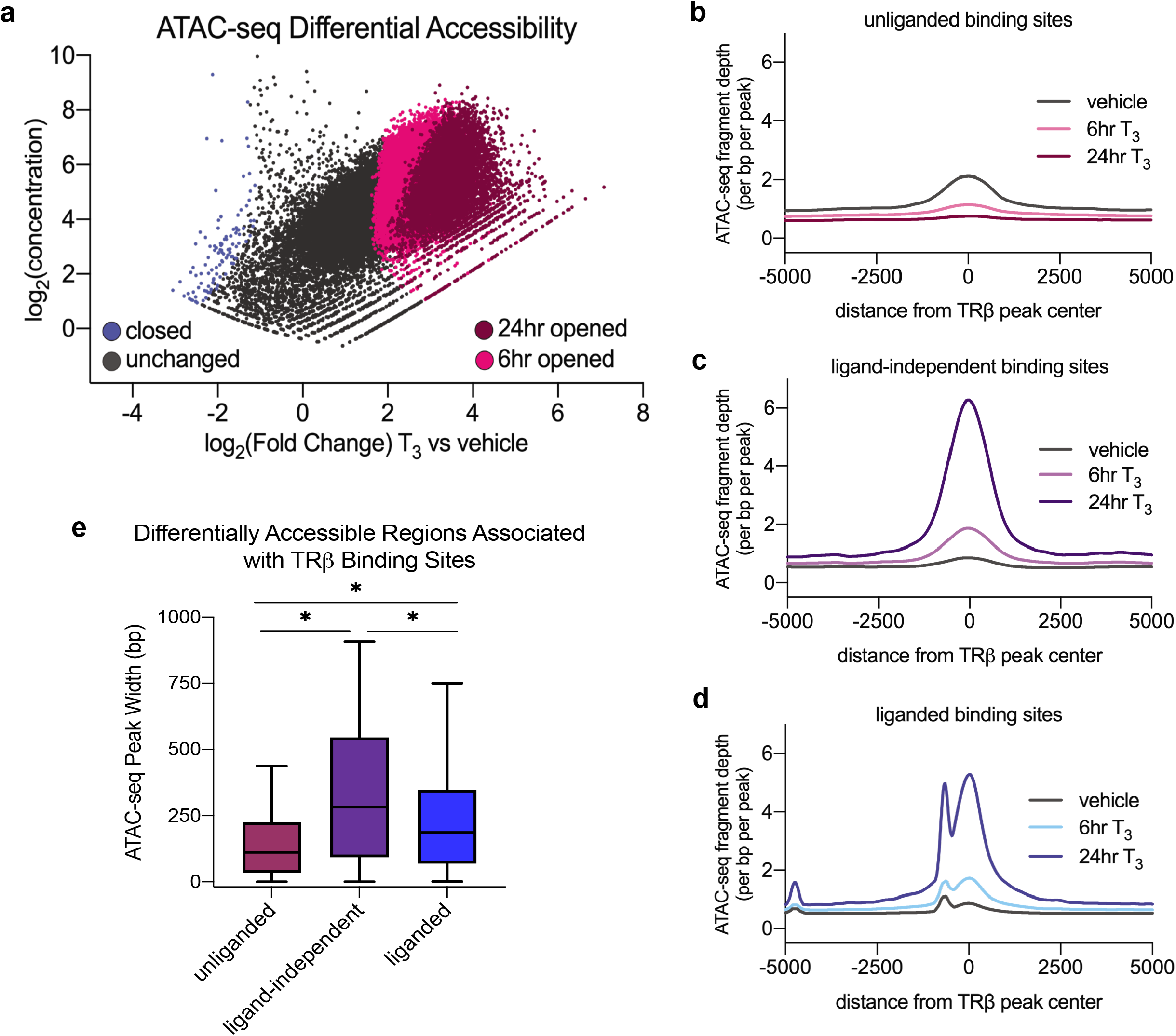
T_3_ induces changes in chromatin accessibility. A. Scatter plot highlights differentially accessible ATAC-seq peaks after 6 and 24 hours of T_3_ treatment. T_3_ induced accessibilty at 7,754 peaks after 6 hours (light pink) and an additional 21,678 peaks after 24 hours (dark pink). 107 peaks were repressed by T_3_ (blue). Differentially accessible peaks are defined as having a log_2_ FC ≥ 1 or ≤ -1 and an FDR ≤ 0.05. ATAC-seq tag desnity is plotted near unliganded (B), ligand-independent (C), and liganded (D) TRβ CUT&RUN peaks. E. Distriubution of peak width of differentially accessible ATAC-seq peaks within 5kb of TRβ binding sites. Satistical significance was determined by one-way ANOVA followed by multiple comparisons; * indicates p < 0.001.

### T_3_ induces changes in the TRβ interactome

To detect protein-protein interactions between TRβ and potential coregulators, including transient cofactors that are exchanged in a T_3_-dependent fashion, we used a live cell proximity-dependent biotin labeling assay (**Figure 1A**). Nthy-ORI cells were transfected with vectors that express a TRβ-miniTurboID fusion construct that can rapidly biotinylate proximal proteins, which were isolated via biotin-affinity purification and subsequently identified by mass spectrometry (**Figure 3A, Supplemental Figure 4**). We identified a total of 1,328 high-confidence proteins that interact with TRβ either in the presence and absence of T_3_. Differential enrichment analysis revealed that 75 of these were gained in the presence of T_3_, and 70 were lost in the presence of T_3_ (**Figure 3B, Supplemental Table 1**). The largest group are unchanged interactions (1,183 proteins). Pathway analysis showed an enrichment of distinctive biological processes associated with the different groups of TRβ interaction partners that change with T_3_ treatment (**Figure 3D**). Interacting proteins that were lost with T_3_ treatment were enriched with transcription repressor activity, demethylase activity, and translation factor activity, while gained proteins were enriched with transcriptional coactivator and RNA polymerase binding activity. Interacting proteins that remained stable were classified as nucleosome remodeling, ATPase activity, and histone binding proteins.

**Figure 3.**
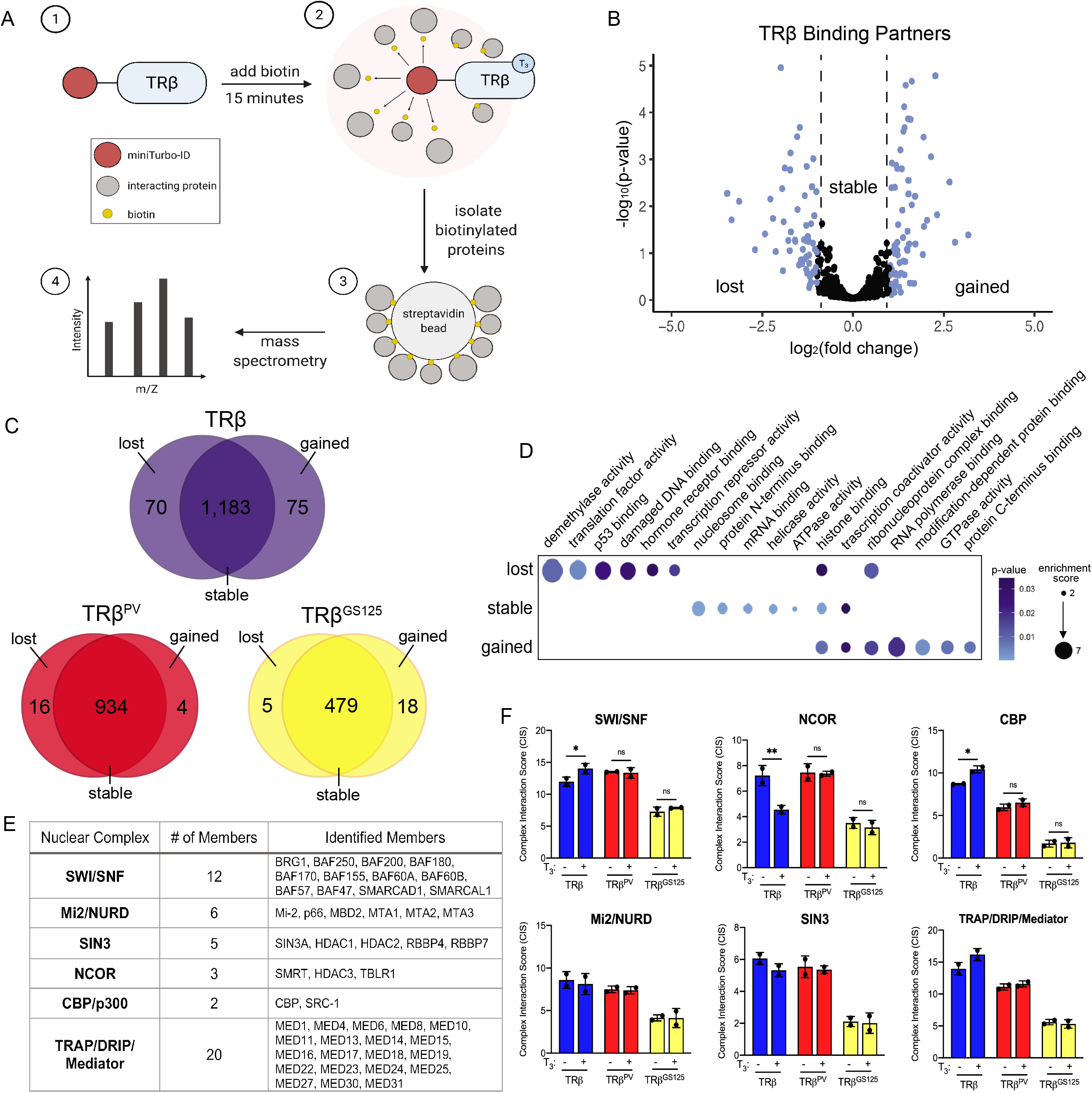
The TRJ3 interactome is altered by T_3_ binding. A. Schematic of miniTurboID proximity labeling assay used to identify TRβ-binding partners. B. Volcano plot depicts differentially enriched TRβ binding partners (DEPs) defined as gained with T_3_ treatment (75), lost with T_3_ treatment (70), or stable (1,183). Differentially enriched proteins are defined as having a log_2_FC ≥ 1 or ≤ -1 and a p-value ≤ 0.05. C. Venn diagrams show DEPs that interact with wildtype TRβ and with TRβ^PV^ (ligand-binding domain mutant), and TRβ^GS125^ (DNA-binding domain mutant). D. GO Molecular Function enrichment of gained, lost, and stable TRβ binding partners. E. Multisubunit chromatin remodeling complexes found to interact with TRβ. F. Complex interaction scores of chromatin remodeling complexes.

To examine the impact of ligand-dependent interactions and DNA-binding dependent interactions, we compared the interaction profiles of a ligand-binding domain mutant (TRβ_PV_)^16^ and a DNA-binding domain mutant (TRβ^GS125^)^17^ with that of wildtype TRβ. The subset of 143 T_3_-induced differential interactions with wildtype TRβ were altered by TRβ^PV^ and TRβ^GS125^ mutants (**Figure 3C**). TRβ^PV^ mutant retained many of the wildtype interactions, but did not exhibit differential binding upon T_3_ treatment, while TRβ^GS125^ mutant lost many interactions entirely. Our analysis of the TRβ interactome is consistent with a co-regulatory shift model^8^, where, rather than an all-or-nothing switch, T_3_ alters the ratio of corepressor to coactivator binding partners.

We focused our analysis on interacting proteins that were likely to participate in chromatin remodeling. Several multi-subunit chromatin remodeling complexes were identified in our proximity labeling assay such as SWI/SNF, Mi-2/NURD, the NCOR and SIN3 co-repressors, and the CBP/p300 and TRAP/DRIP/Mediator co-activators (**Figure 3E**). Each of these has been previously linked to TRβ gene regulation^18-22^. We identified several members of each of these complexes as in our proteomic dataset. An interaction score was calculated for TRβ-associated chromatin remodeling complexes by dividing the sum of the signal intensity of each subunit in the complex by the total subunits identified (**Figure 3F**). The Mi-2/NURD, SIN3, and TRAP/DRIP/Mediator complexes did not have significant changes in their interaction scores. There was an increase in the CBP coactivator complex interaction score and decrease in the NCOR corepressor complex interaction score with T_3_, consistent with previous studies of their interaction with TRβ^18^. The SWI/SNF complex also significantly increased with T_3_. TRβ^PV^ and TRβ^GS125^ mutations both inhibited these T_3_-dependent interactions.

### TRβ differentially recruits SWI/SNF complexes

Upon closer examination of the SWI/SNF complex subunits that were identified as TRβ interacting proteins, it became apparent that TRβ interacts with two different subspecies of SWI/SNF complexes: canonical BAF and polybromo-associated BAF (PBAF). Each of these contains one of the mutually exclusive ATPase catalytic subunits, (BRG1 or BRM) which are necessary for nucleosome displacement, array of accessory components, and a few subspecies-specific subunits^23^ (**Figure 4A**). Our proteomic analysis identified the BRG1 core subunit, many of the accessory components, and three subspecies-specific subunits.

**Figure 4.**
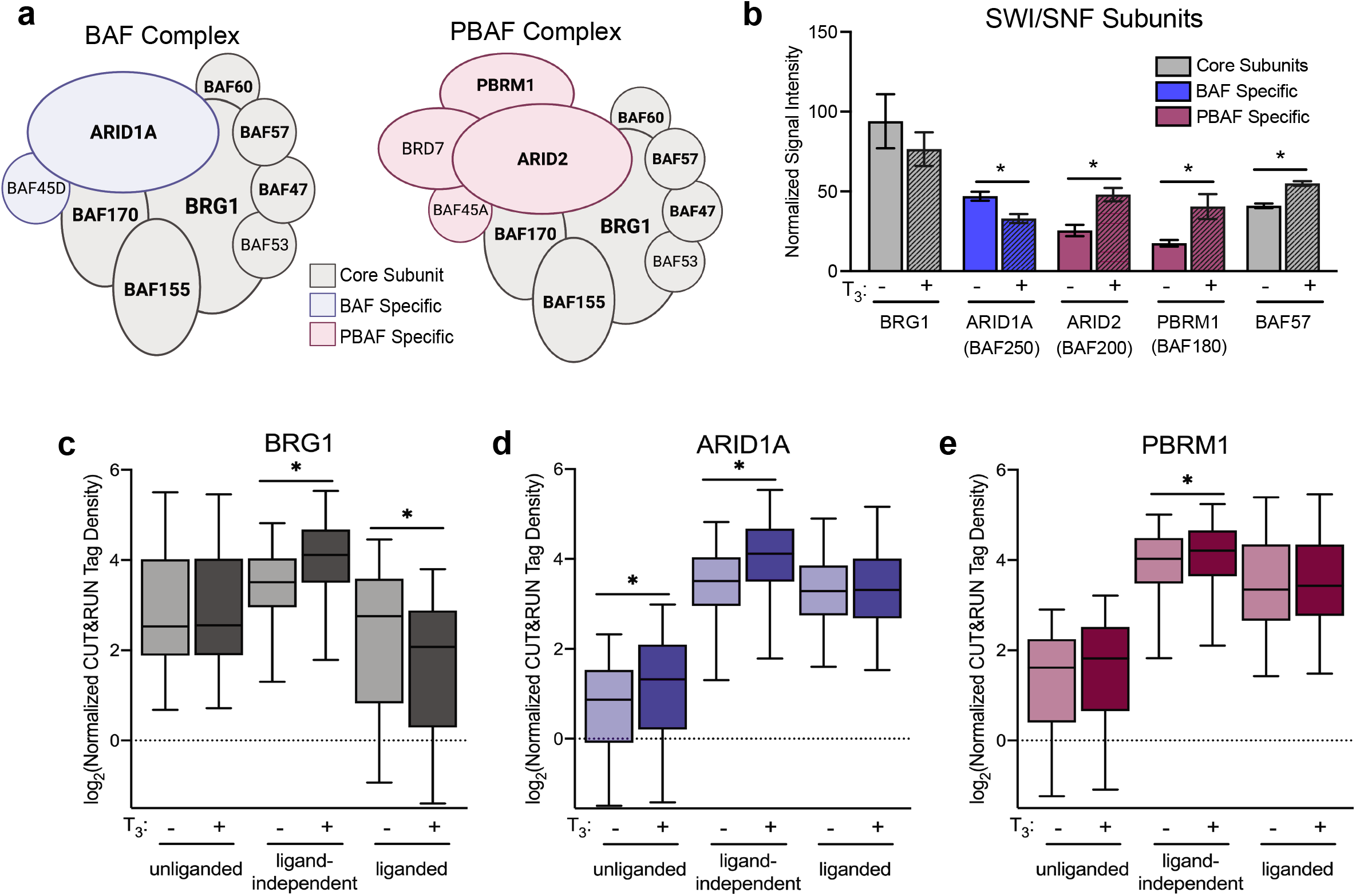
BAF and PBAF complexes differentially interact with TRβ. A. Diagram of BAF and PBAF complex subunits. Core subunits are colored in grey, BAF specific subunits are colored in blue, and PBAF specific subunits are colored in pink. Labels of subunits identified as TRβ binding partners by miniTurboID proximity-labeling assay are bolded. B. Signal intensity of differentially enriched SWI/SNF subunits identified in miniTurboID proximity-labeling assay in the presence and absence of T_3_. Significance (* p<0.05) was determined by paired t-test. Box plots demonstrate differential CUT&RUN tag density of BRG1 (C), ARID1A (D), and PBRM1 (E) subunits at TRβ binding sites. Significance (* p<0.05) was determined by paired t-test.

Enrichment of the BAF-specific subunit ARID1A (BAF250) was decreased with T_3_ treatment, while the PBAF-specific subunits ARID2 (BAF200) and PBRM1(BAF180) were increased in the presence of T_3_ (**Figure 4B**). BAF57 enrichment was also increased in the presence of T_3_. BRG1 was not differentially enriched.

Based on differential enrichment observed in our proximity ligation data, we performed CUT&RUN to determine T_3_-induced changes in genomic binding of ARID1A as a representative of BAF complexes, PBRM1 as a representative of PBAF complexes, as well as BRG1 (**Supplemental Figure 1B-D**). We then examined the binding of each of these factors specifically at TRβ binding sites. BRG1 CUT&RUN tag density was unchanged with T_3_ at unliganded sites, while ARID1A tag density was low with a slight increase with T_3_, and PBRM1 tag density was low (**Figure 4C**). This indicates that unliganded TRβ may recruit BRG1 to prime its binding sites in a manner similar to a mechanism described for the glucocorticoid receptor^24^. Ligand-independent sites had an increase in all three SWI/SNF components with T_3_, indicating that both BAF and PBAF complexes are recruited (**Figure 4D**). Liganded binding sites had an increase BRG1 tag density, while ARID1A and PBRM1 remained stable (**Figure 4E**).

To further examine differential recruitment of BAF and PBAF complexes, TRβ peaks were annotated based on whether they occurred near a promoter (< 5kb from nearest TSS) or in a distal regulatory region (> 5kb from nearest TSS). The CUT&RUN tag density of SWI/SNF subunits were quantified near two subsets of peaks (**Figure 5A**). As expected, BRG1 was recruited to both sites in the presence and absence of T_3_. ARID1A was recruited in a T_3_-dependent manner to both TRβ-bound promoters and distal binding sites. PBRM1, however, was recruited preferentially to TRβ-bound promoters and not to distal binding sites. Visualization of representative binding sites in genome browser shots (**Figure 5D**) further demonstrated differential recruitment. There is a much greater degree of change in chromatin accessibility at 6 and 24 hours near TRβ-bound promoters (**Figure 5B**) then at distal binding sites (**Figure 5C**). Given that both BAF and PBAF complexes are recruited to promoters, both complexes may be required to facilitate the large changes in chromatin accessibility observed (**Figure 5B**).

**Figure 5.**
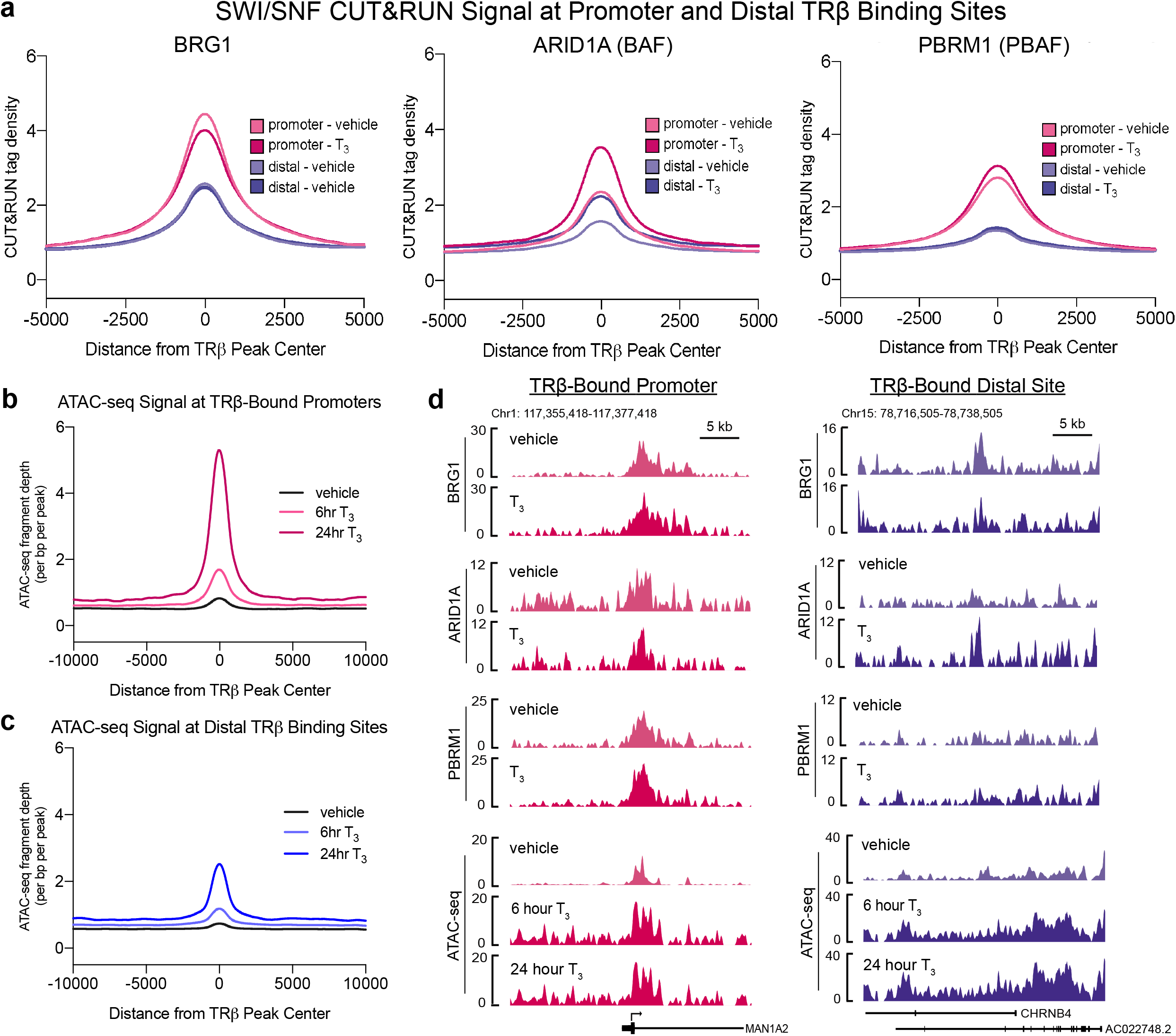
BAF and PBAF complexes are differentially recruited to TRβ binding sites. A. CUT&RUN tag density of BRG1, ARID1A (BAF-specific), and PBRM1 (PBAF-specific) near TRβ binding sites classified as promoters (< 5kb from nearest TSS) or distal binding sites (> 5kb from nearest TSS). BRG1 and ARID1A are recruited to promoter and distal sites, PBRM1 is preferentially recruited to promoter sites. B. ATAC-seq tag density is increased upon T_3_ treatment for 6 and 24 hours near promoter binding sites. C. ATAC-seq tag density is increased upon T_3_ treatment for 6 and 24 hours near distal binding sites. D. Genome browser shots (fold enrichment over IgG) highlight differential recruitment of SWI/SNF complexes to a representative promoter and distal binding site.

### TRβ chromatin interactions are correlated with target gene expression

To determine the effect of T_3_ treatment on gene expression, Nthy-ORI cells were treated with 10nM T_3_ or vehicle control for 6 and 24 hours and global transcriptomic analysis by RNA-seq was performed. Differential gene expression analysis was performed comparing each treatment to the corresponding control. We determined that 366 and 368 genes were up and downregulated, respectively, by T_3_ at 6 hours (**Figure 6A, Supplemental Table 2**). The effect was increased at the 24-hour period where 480 and 667 up and downregulated genes, respectively. Enriched biological function pathways within early and late DEGs were compared using Ingenuity Pathway Analysis software (**Figure 6C**). Highly enriched upregulated pathways included cellular homeostasis, survival, viability, and cell cycle progression. Transcription and protein synthesis switched from down- to upregulated between early and late T_3_ treatment, while carbohydrate metabolism switched from up- to downregulated. Apoptosis, cell transformation, and ER stress response were downregulated. BETA^25^ was used to predict whether T_3_-regulated genes are likely to be direct transcriptional targets of TRβ based on proximity of a TRβ peak to the TSS (**Figure 6B**). 63% (230/366) of early upregulated genes and 51% (245/480) of late upregulated genes were predicted to be direct targets of TRβ. This suggests that increases in chromatin accessibility that occur in the time between early and late T_3_ timepoints (**Figure 2A**) may allow TRβ to access additional binding sites for induction of a subsequent set of target genes. Conversely, 48% (177/368) of early and 18% (120/667) of late downregulated genes were predicted to be direct targets of TRβ, which indicates that direct TRβ-induced gene repression occurs more immediately and the additional downregulation we observed may be secondary effects downstream of the early transcriptomic effects of T_3_-treatment.

**Figure 6.**
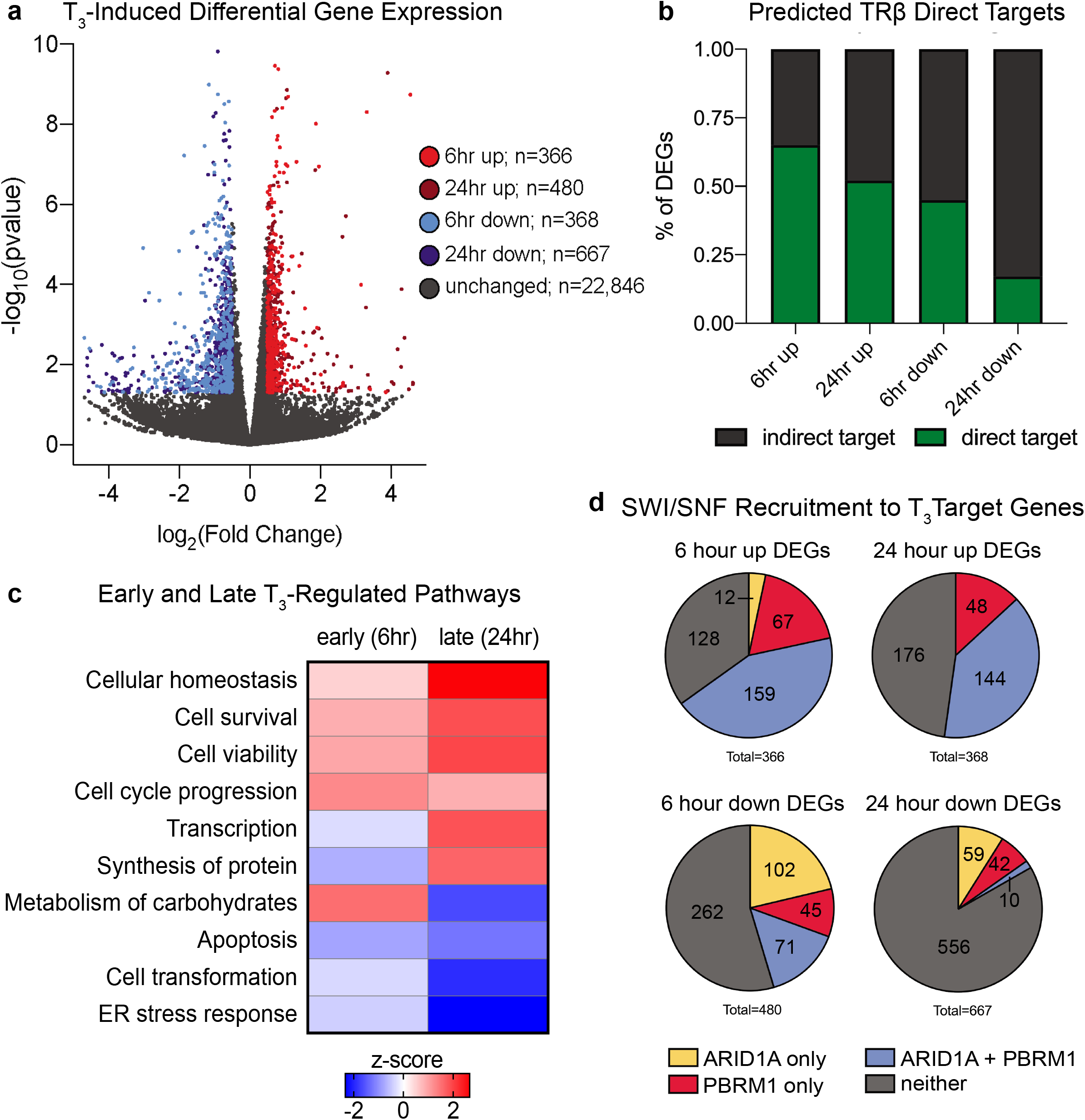
TRβ chromatin interactions are correlated with target gene expression. A. T_3_ -induced and -repressed genes determined by RNA-seq after 6 and 24 hours of T_3_ treatment in two independent biological replicates. Differentially expressed genes (DEGs) are defined as having a log2FC ≥ 0.5 or ≤ -0.5 and an p-value ≤ 0.05. B. BETA regulatory potential prediction of TRβ direct targets among DEGs after 6 and 24 hours of T_3_ treatment. Direct targets have a log2FC ≥ 0.5 or ≤ -0.5 p-value ≤ 0.05 and a TRβ peak within 10kb of the TSS. C. IPA analysis reveals differentially enriched biological functions after early and late T_3_ treatment. D. Proportion of DEGs with ARID1A (yellow) or PBRM (red) binding exclusively both ARID1A and PBRM1 (blue) or neither within 10kb of the promoter.

We examined SWI/SNF complex recruitment to promoters of T_3_-induced differentially expressed genes (**Figure 6D**). A majority of early upregulated genes have a SWI/SNF complex binding site within 10kb of the TSS; most have both ARIDIA and PBRM1 peaks, while a minority had exclusively one or the other. There was a similar trend in the late upregulated genes. In contrast, a majority of downregulated genes, both early and late, did not have SWI/SNF binding near their promoter. However, among those that did there was a preference for ARID1A recruitment over PBRM1. Combined, these results suggest that SWI/SNF complexes participate in T_3_-induced gene regulation near promoters of target genes, particularly in the context of upregulation. For upregulation of gene expression, both BAF and PBAF complexes are likely to be recruited to remodel the proximal promoter region, while BAF complexes may be preferred for remodeling of downregulated promoters.

## DISCUSSION

Although it is now appreciated that the classic model for TRβ interaction with chromatin is oversimplified, a consensus has yet to be reached on an updated model. Based on the data presented here, we propose a multi-modal regulation model where TRβ has at least three distinct modes of binding and remodeling chromatin (**Figure 7**). In agreement with previous genome-wide studies^4-6,8^, we observed significant shifts in TRβ occupancy in the presence and absence of T_3_. Unliganded TRβ binds to chromatin with limited effects on chromatin accessibility, but much of this binding is lost upon T_3_ treatment. Liganded binding sites represent the vast majority of TRβ binding, and they are associated with significant changes in localized chromatin accessibility (**Figure 1A,B**; **Figure 2C**). Intriguingly, liganded binding sites show a clear increase in enrichment upon the addition of T_3_, however most sites have some, albeit low, enrichment before T_3_ is added (**Figure 1B**). This suggests that TRβ interactions with this type of site are transient and are stabilized by ligand binding and recruitment of coregulators, consistent with a dynamic assisted loading model^15,26^. Since they are numerous and broadly distributed across the genome, there are likely multiple functions of liganded binding sites that further studies may clarify. While some of these binding sites occur near proximal promoters and facilitate direct induction or repression of gene expression, many occur in distal regulatory elements and may regulate enhancers or coordinate higher order chromatin structure. Ligand-independent binding sites are defined by enrichment both in the presence and absence of T_3_, and substantial induction of chromatin accessibility (**Figure 1A,B; Figure 2D**). While they have a clear functional importance for regulation of gene expression, these binding sites represent a small minority of the overall cistrome of TRβ. Notably, ligand-independent binding sites have many of the characteristics originally described in the bimodal switch model such as their proximity to transcriptional start sites, high frequency of full-length DR4 TREs, and high regulatory potential.

**Figure 7.**
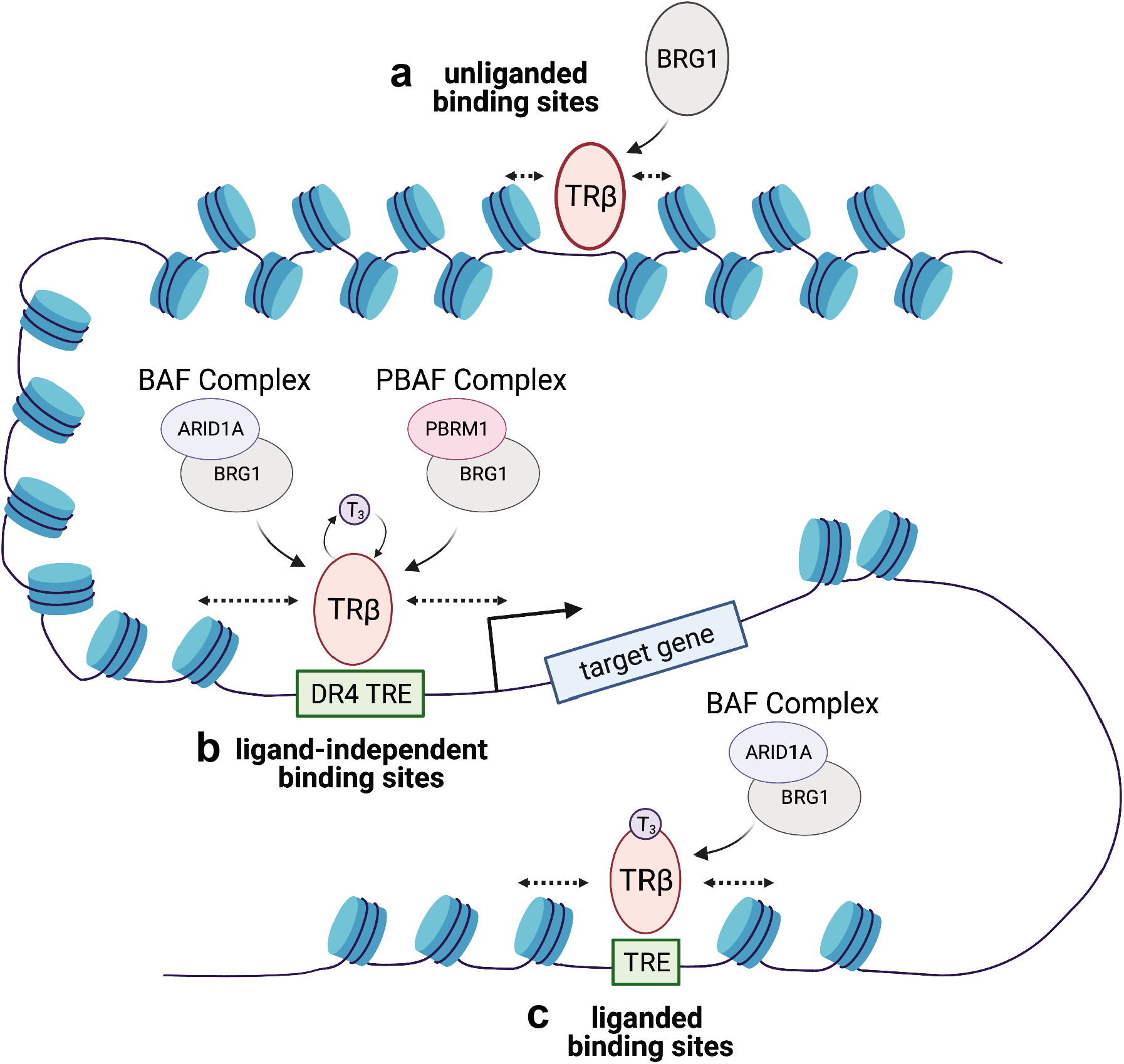
Multi-modal regulation model for TRJ. chromatin interaction and recruitment of SWI/SNF complexes. A. BRG1 is recruited to unliganded TRβ binding sites for modest changes in chromatin accessibility. B. BAF and PBAF complexes are both recruited to ligand-independent TRβ binding sites near the TSS of target genes to facilitate broad changes in chromatin accessibility. C. BAF complexes are specifically recruited to liganded TRβ binding sites to facilitate T_3_ -induced changes in chromatin accessibility at distal regulatory regions.

TRβ binding sites are also characterized by differential recruitment of SWI/SNF chromatin remodelers. BRG1 alone is recruited to unliganded and liganded sites in the absence of T_3_ (**Figure 4D**) suggesting that that it may be recruited to prime binding sites for TRβ occupancy, a mechanism which has been clearly delineated for the glucocorticoid receptor^24^. BAF and PBAF complexes, each with unique subunits which dictate their precise function, are recruited to TRβ binding sites within promoters and likely contribute directly to the changes in chromatin accessibility and recruitment of other transcription factors to alter target gene expression. Both BAF and PBAF complexes have been implicated previously in hormone-dependent gene regulation^27,28^, however the distinct functional role of each when they are recruited by a nuclear receptor to the same location remains unclear. BAF complexes are preferentially recruited by TRβ to binding sites that occur in distal regulatory regions. This might be a mechanism by which TRβ primes these binding sites or organizes higher order chromatin structure to promote persistent changes in gene expression. These multifaceted interactions with a single chromatin remodeling complex illustrate the importance of the proposed coregulator shift model^8^ over a complete coregulator switch.

As a deeper understanding of the variety of ways in which TRs regulate gene expression and coordinate a network of cofactors is developed, it is important that the models we use reflect their multidimensional function. We suggest a model for multi-modal regulation by TRβ that has at least three distinct modes defined by their T_3_-dependent occupancy, changes in accessibility, and differential recruitment of chromatin remodelers. Collectively, this study provides a next-generation model for TRβ interactions with chromatin, and lays a foundation for further studies of TRβ regulation of gene expression and recruitment of key cofactors in both normal cells and in disease models.

## METHODS

### Cell Culture and Hormone Treatments

Nthy-ORI cells (Sigma) were routinely cultured in RPMI 1640 growth media with L-glutamine (300 mg/L) (Sigma), sodium pyruvate and nonessential amino acids (1%) (Cellgro/Mediatech), supplemented with 10% fetal bovine serum (Peak Serum) and penicillin-streptomycin (200 IU/L) (Cellgro/Mediatech) at 37°C, 5% CO_2_, and 100% humidity. For T_3_ treatments, cells were hormone-starved for 24 hours in growth media substituted with phenol-red free RPMI 1640 and charcoal-stripped fetal bovine serum (Sigma) prior to the addition of 10nM T_3_ or NaOH vehicle for the indicated time course. Nthy-ORI cell line was authenticated by the Vermont Integrative Genomics Resource at the University of Vermont with short tandem repeat profiles using the Promega GenePrint10 platform.

### CUT&RUN

#### Sample collection and Sequencing

CUT&RUN was performed as described ^13^. Briefly, Nthy-ORI cells were harvested, washed, and bound to activated Concanavalin A coated magnetic beads (Epicypher 21-1401). Cells were then permeabilized with Wash buffer (20mM HEPES pH 7.5, 150mM NaCl, 0.5 mM spermidine 0.05% digitonin). Permeabilized cells were then incubated with the indicated antibody (Supplemental Table 3) at 4°C with constant agitation overnight. Cells were washed twice more before incubation with recombinant p-AG MNase (Epicypher 15-1016) at 4°C for 2 hours. Liberated DNA was purified, and libraries were prepared using the NEB Ultra FS II DNA Library Kit (NEB E6177) and amplified with 14 cycles of PCR. Amplified libraries were then purified with AMPure beads (Agencourt), quantified via Qubit (Life Technologies), and quality was assessed using the BioAnalyzer (Agilent) High-Sensitivity DNA kit. CUT&RUN libraries were pooled and sequenced on the Illumina HiSeq 1500 with 100 bp paired-end reads.

#### Data Analysis

Quality scores across sequenced reads were assessed using FASTQC. Illumina adapters were removed using Trim-Galore. Paired-end reads were mapped to hg38 using Bowtie2, and peaks were called using MACS2. Consensus peak sets for downstream analysis were derived using IDR ^29^ using two replicates (Supplementary Figure 2A) per target and a cut-off of 0.05.

### ATAC-seq

#### Sample collection and Sequencing

ATAC-Seq was performed as previously described ^30^ using 50,000 Nthy-ORI cells with two biological replicates per condition. Libraries were generated using custom Nextera barcoded primers ^30^ and were amplified by PCR for a total of 10 cycles. Amplified libraries were then purified with AMPure beads (Agencourt), quantified using a Qubit (Life Technologies), and quality was assessed using the BioAnalyzer (Agilent) High-Sensitivity DNA kit. ATAC-seq libraries were then pooled and sequenced on the Illumina HiSeq 1500 with 100 bp paired-end reads.

#### Data Analysis

Quality scores across sequenced reads were assessed using FASTQC. Nextera adapters were removed using Trim-Galore. Paired-end reads were mapped to hg38 using Bowtie2, and peaks were called using MACS2. DiffBind^31^ was used to identify regions of differential accessibility. Consensus peak sets for downstream analysis were derived using IDR ^29^ using two replicates (Supplementary Figure 2) per target and a cut-off of 0.05.

### RNA-seq

#### Sample collection and Sequencing

Nthy-ORI cells were treated 10nM T_3_ or vehicle for 6 or 24 hours prior to sample collection. Total RNA was extracted and purified using RNeasy Plus Kit (Qiagen) according to manufacturer’s protocol. This was repeated to collect a total of three biological replicates per condition. Purity of the total RNA samples was assessed via BioAnalyzer (Agilent) and samples with an RNA integrity score >8 were used for library construction. rRNA was depleted from 1 μg of total RNA with the RiboErase kit (KAPA Biosystems). Strand-specific Illumina cDNA libraries were prepared using the KAPA Stranded RNA-Seq library preparation kit with 10 cycles of PCR (KAPA Biosystems). Library quality was assessed by BioAnalyzer (Agilent) to ensure an average library size of 300bp and the absence of excess adaptors in each sample. RNA-Seq libraries were pooled and sequenced on the Illumina HiSeq 1500 with 50 bp single-end reads.

#### Data Analysis

Quality scores across sequenced reads were assessed using FASTQC. All samples were high quality. For alignment and transcript assembly, the sequencing reads were mapped to hg38 using STAR. Sorted reads were counted using HTSeq and differential expression analysis was performed using DESeq2. Genes with a p-value of <0.05 and a log_2_ fold change greater than 1 or less than -1 were considered differentially expressed (Supplemental Table 2).

### Proximity Labeling by miniTurboID

#### Cloning of 3xHA-miniTurbo-TRβ

3xHA-miniTurbo-NLS_pCDNA3 vector was a gift from Dr. Alice Ting (Addgene plasmid # 107172; http://n2t.net/addgene:107172 ; RRID: Addgene_107172)^32^. 3xHA-miniTurbo-NLS_pCDNA3 was linearized by PCR and THRB cDNA was inserted via HiFi Assembly Cloning (NEB E5520S) to create the 3X-HA-miniTurbo-TRβ vector. Site directed mutagenesis (NEB E00554) was used to create mutant 3X-HA-miniTurbo-TRβ-GS^125^ and 3X-HA-miniTurbo-TRβ-PV vectors. Successful insertion of the THRB cDNA and site-directed mutagenesis were confirmed by Sanger sequencing. Expression of fusion constructs from the cloned vectors was confirmed by Western blot (Supplemental Figure 3A).

#### Transfection and Biotin Labeling

Nthy-ORI cells were grown as a monolayer in DMEM-F12 (Cellgro/Mediatech) supplemented with 10% fetal bovine serum and penicillin-streptomycin (200 IU/L) in 15cm cell culture dishes. Cells were transfected at approximately 80% confluency with 20μg of plasmid DNA using 25μL Lipofectamine 3000 for 24 hours. BioID samples were simultaneously labeled using 250μM biotin and treated with 10nM T_3_ or vehicle for 15 minutes. Labeling was stopped by placing cells on ice and washing three times with ice-cold PBS. Cells were detached from the plate and collected by centrifugation. The cell pellet was subjected to nuclear protein extraction using the NE-PER Protein Extraction Kit (ThermoFisher 78833) with the addition of Protease Inhibitor Cocktail (Thermo Scientific 781410) per the manufacturer’s instructions.

#### Sample Preparation and Mass Spectrometry

To enrich biotinylated proteins, 1mg of nuclear extract was incubated for 30 minutes rotating at room temperature with 100uL of streptavidin-coated magnetic beads (Invitrogen 65001). The supernatant was removed, and the beads were washed three times with high salt RIPA buffer (100mM Tris pH 9.0, 500mM LiCl, 150mM NaCl, 1% Igepal/NP-40, 1% deoxycholic acid). Washed beads were then resuspended in Laemmli sample buffer and boiled for 10 minutes to denature and release the biotinylated proteins from the beads. The eluents were loaded onto 10% Tris-Glycine gels (Invitrogen XP00100BOX), and separated by SDS-PAGE. Gels were then silver stained (Pierce 24600) prior to band excision for mass spectrometry (Supplemental Figure 3B). LC-MS was performed using an LTQ-Orbitrap instrument (ThermoFisher).

#### Data Analysis

Data acquired by mass spectrometry was quantified using MaxQuant label-free quantification (LFQ) workflow, and LFQ values were used to calculate differential enrichment of identified proteins between experimental conditions using the DEP Bioconductor package^33^ (Supplemental Figure 3C, Supplemental Table 1). Proteins with a p-value of <0.05 and a log_2_ fold change greater than 1 or less than -1 were considered differentially enriched. Proteins found to be enriched in the empty vector control group were excluded from wildtype and mutant TRβ datasets and were not used for downstream analysis.

## Data Availability

All raw and processed next generation sequencing data associated with this manuscript is available for download from the NCBI Gene Expression Omnibus repository under accession code GSE168954.

## Acknowledgements

The research reported here was supported by grants from National Institutes of Health U54 GM115516 for the Northern New England Clinical and Translational Research Network; National Cancer Institute 1F99CA245796-01; and UVM Larner College of Medicine. Next-generation sequencing was performed in the Vermont Integrative Genomics Resource Massively Parallel Sequencing Facility supported by the University of Vermont Cancer Center and the UVM Larner College of Medicine. Mass spectrometry was performed at the Vermont Biomedical Research Network Proteomics Core Facility with support from Dr. Bin Deng. Figures 3A, 4A, and 7 were created with Biorender. The authors would like to thank Dr. Jane Lian for her thoughtful discussions and guidance on this project.

## Author Contributions

NEG, SF, and FEC conceptualized and designed this study; NEG and JAT performed the experiments and prepared the sequencing libraries. NEG analyzed the data with help from JRB, and SF. NEG drafted the manuscript and it was edited by JAT, JRB, SF, and FEC.

### Conflict of Interest Statement

The authors declare that they have no conflict of interest.

## Notes

### Competing Interest Statement

The authors have declared no competing interest.

