## Supplemental Figures 1-4 for "Thyroid Hormone Dependent Transcriptional Programming by TRβ Requires SWI/SNF Chromatin Remodelers"

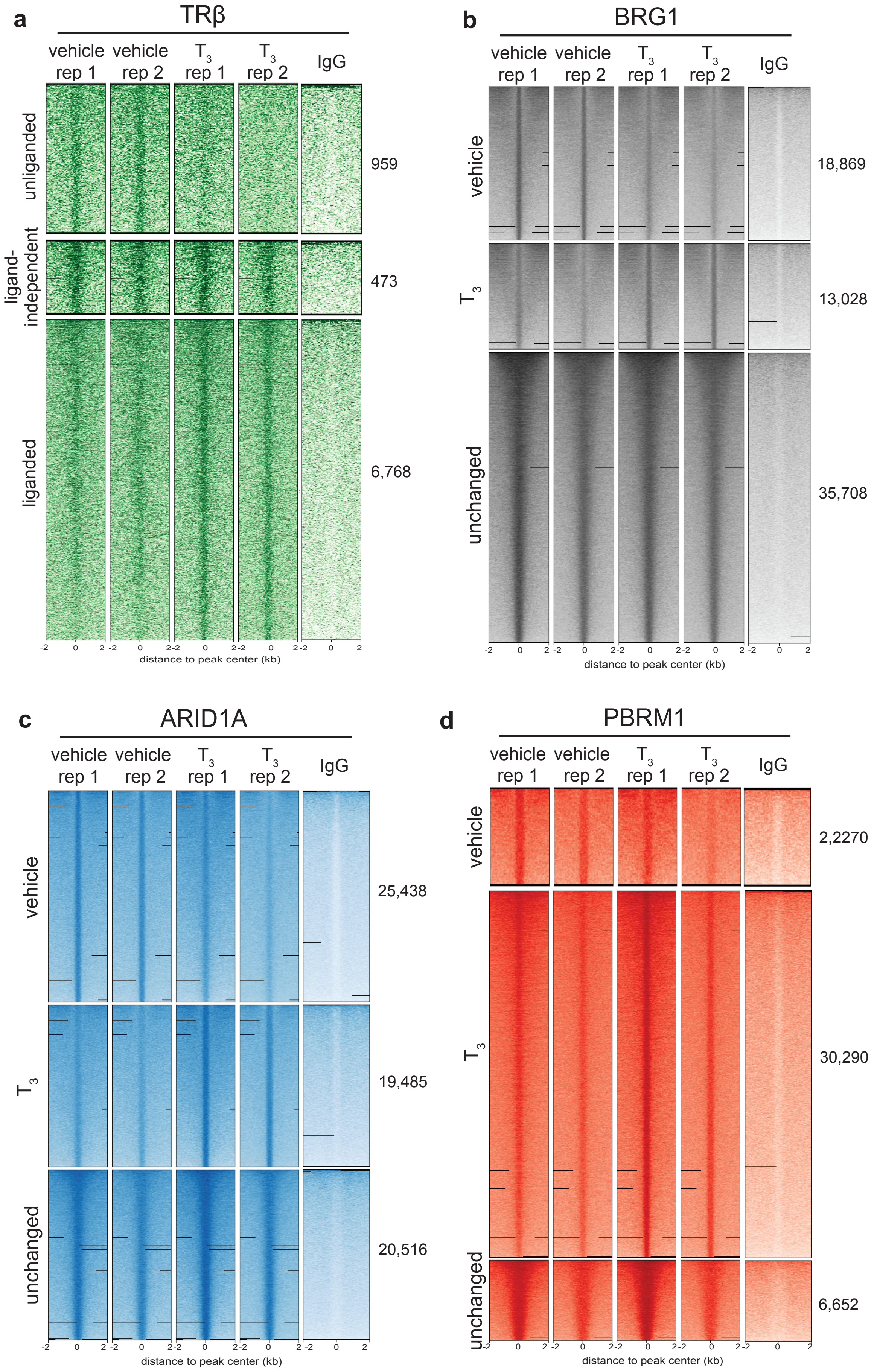


**Supplemental Figure 1.** CUT&RUN signal at TRβ (A), BRG1 (B), ARID1A (C), and PBRM1 (D) binding sites in the presence and absense of $T_3$ demonstrates concordance between replicates and enrichment compared to IgG control.

a. <u>DR4 TRE:</u> 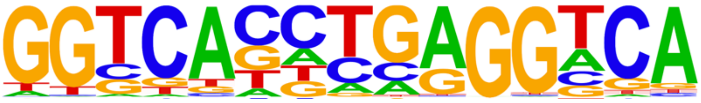

| Type of TRβ Binding Site | p-value | % of peaks with motif |
|---|---|---|
| unliganded | 1e-2 | 10.4% |
| ligand-independent | 1e-36 | 15.4% |
| liganded | 1e-10 | 61.6% |

b. <u>Half site TRE:</u> 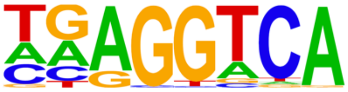

| Type of TRβ Binding Site | p-value | % of peaks with motif |
|---|---|---|
| unliganded | 1e-8 | 20.5% |
| ligand-independent | 1e-18 | 41.6% |
| liganded | 1e-20 | 32.9% |

**Supplemental Figure 2.** Transcription factor motif analysis revealed significant enrichment of full length direct-repeat (DR4) thyroid hormone receptor response elements (TRE) (A) and TRE half sites (B) in all three types of TRβ binding sites.

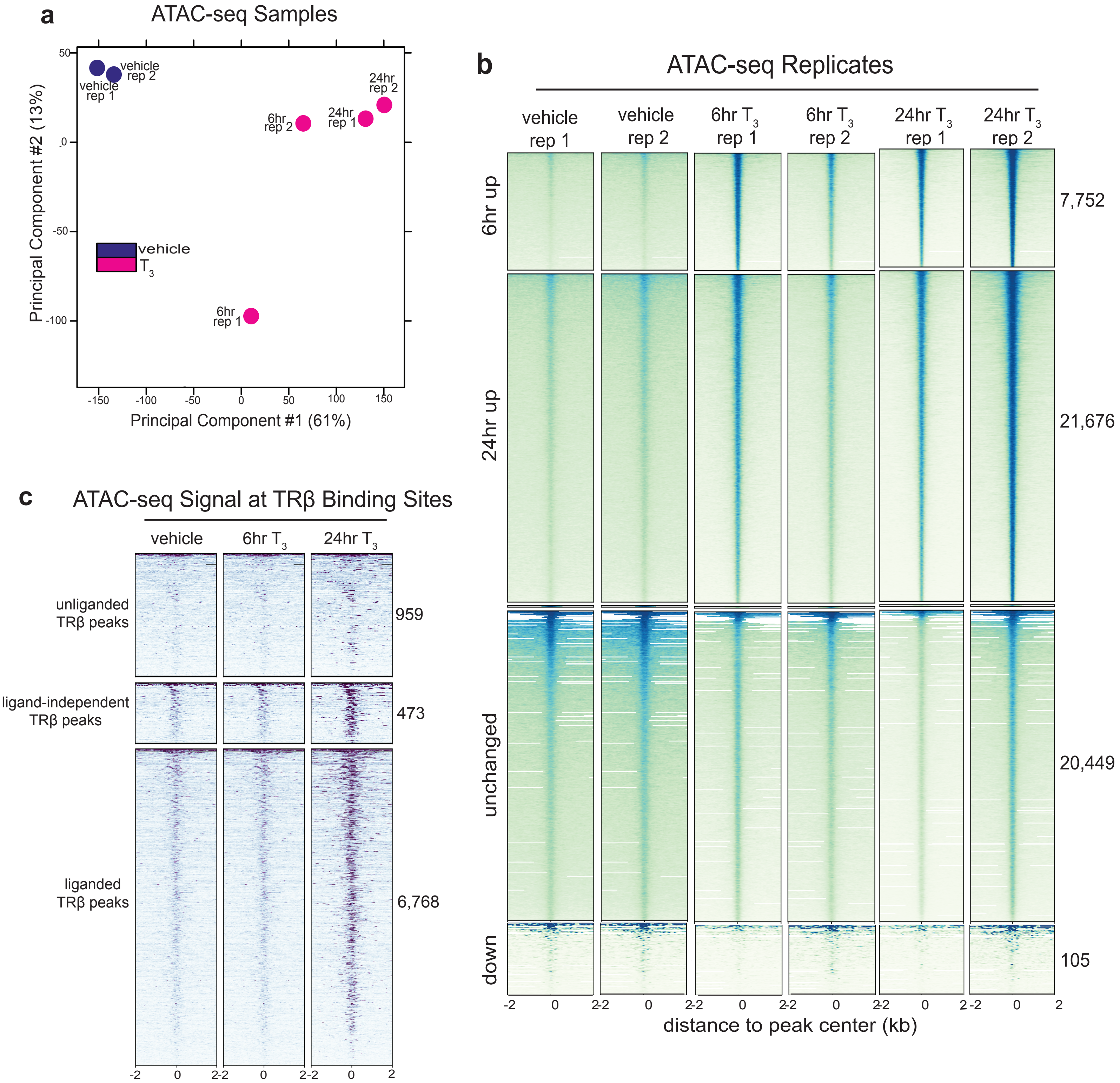


**Supplemental Figure 3.** A. PCA plot demonstrates concordance between replicate ATAC-seq profiles and separation between samples treated with $T_3$ compared to vehicle. B. ATAC-seq signal at differentially accessible peaks in the presence and absense of $T_3$. C. ATAC-seq signal at unliganded, liganded, and ligand-idependent TRβ binding sites.

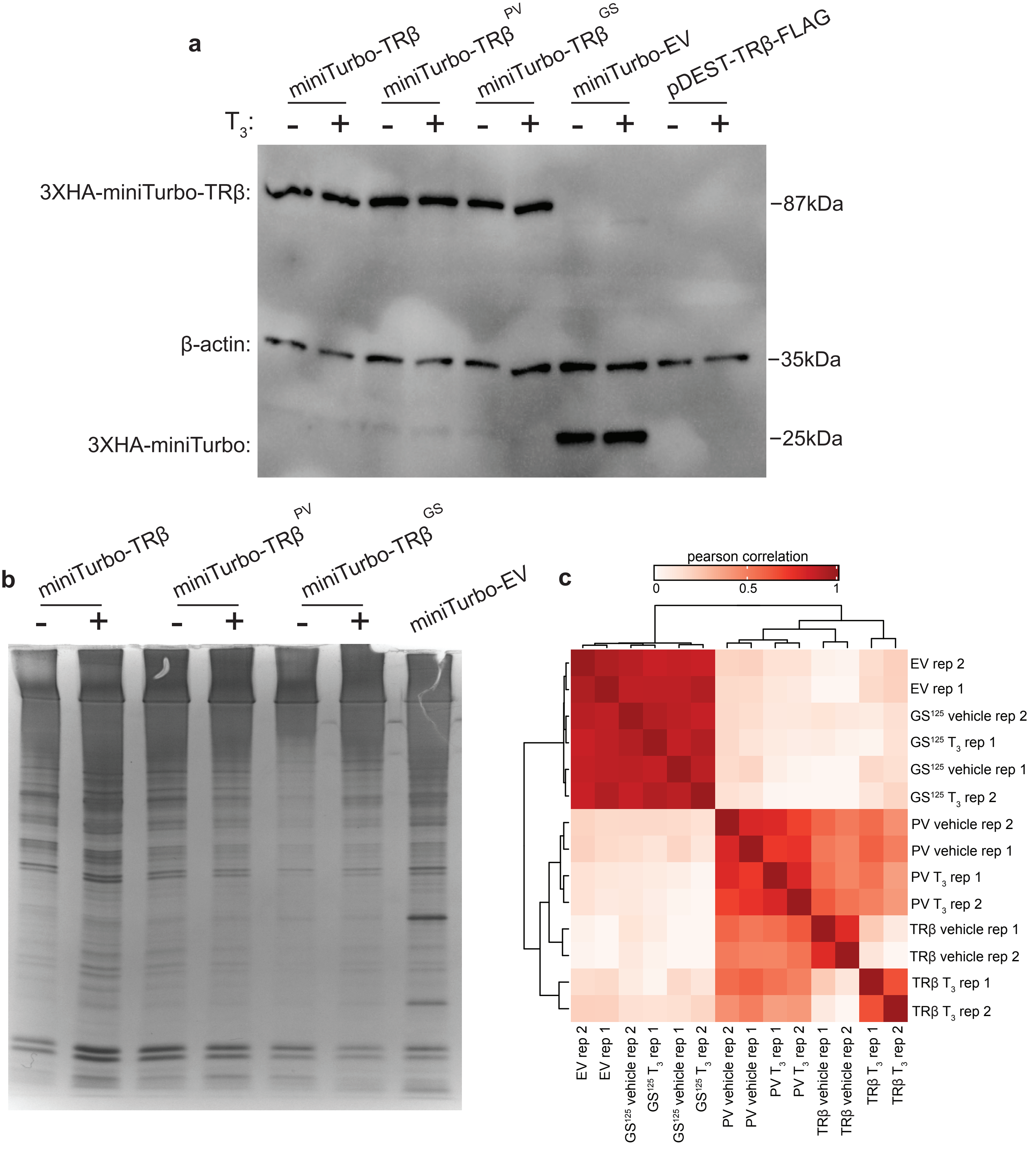


**Supplemental Figure 4.** A. Western blot depicts protein expression levels of miniTurbo-TRβ and miniTurboID-EV constructs after transfection into Nthy-ORI cells. B. Silver stain shows biotinylated proteins isolated by streptavidin bead pull-down from Nthy-ORI cell nuclear lysates collected after transfection with miniTurbo-TRβ and miniTurboID-EV constructs. C. Pearson correlation heatmap demonstrates concordance between replicate protein profiles identified by mass spectrometry.

### Supplemental Table 3. Antibody Table

| Target | Catalog Number | Company | Use |
|---|---|---|---|
| TRβ | ab5622 | abcam | CUT&RUN (1μg) |
| TRβ | MA1-216 | ThermoFisher | Western blot (1:1000) |
| BRG1 | ab110641 | abcam | CUT&RUN (0.5μg), Western blot (1:5000) |
| ARID1A | 12354 | Cell Signaling | CUT&RUN (0.5μg), Western blot (1:1000) |
| PBRM1 | AB_2793612 | Active Motif | CUT&RUN (0.5μg) |
| rabbit IgG | 13-0042 | Epicypher | CUT&RUN (0.5μg) |
| HA tag | 3724 | Cell Signaling | Western blot (1:1000) |
| goat-anti-mouse | 401253 | Calbiochem | Western blot (1:10000) |
| goat-anti-rabbit | 112-035-175 | Jackson | Western blot (1:10000) |
